# Organization of state transitions in the resting-state human cerebral cortex

**DOI:** 10.1101/404418

**Authors:** Jiyoung Kang, Chongwon Pae, Hae-Jeong Park

## Abstract

The resting-state brain is often considered a nonlinear dynamic system transitioning among multiple coexisting stable states. Despite the increasing number of studies on the multistability of the brain system, the processes of state transitions have rarely been systematically explored. Thus, we investigated the state transition processes of the human cerebral cortex system at rest by introducing a graph-theoretic analysis of the state transition network. The energy landscape analysis of brain state occurrences, estimated using the pairwise maximum entropy model for resting-state fMRI data, identified multiple local minima, some of which mediate multi-step transitions toward the global minimum. The state transition among local minima is clustered into two groups according to state transition rates and most inter-group state transitions were mediated by a hub transition state. The distance to the hub transition state determined the path length of the inter-group transition. The cortical system appeared to have redundancy in inter-group transitions when the hub transition state was removed. Such a hub-like organization of transition processes disappeared when the connectivity of the cortical system was altered from the resting-state configuration. In summary, the resting-state cerebral cortex has a well-organized architecture of state transitions among stable states, when evaluated by nonlinear systematic approach.

## INTRODUCTION

A dynamic complex system can possess several stable states (attractors) for a given set of system parameters^1-6^. If a system has multiple coexisting stable states and can switch among them in response to noise or intrinsic perturbations to the system, it is generally referred to as a multistable system^7,8^. In this respect, the brain at rest can be considered as a system showing multistability^1-7,9,10^. A repertoire of independent spatial components^11-13^ and subnetwork components^14^ from spontaneous fluctuations of blood oxygenation level dependent (BOLD) functional magnetic resonance imaging (fMRI) signals are examples that suggest multistability of the resting state brain. From the perspective of the multistable brain, the conventional term “resting state” is not a homogeneous state but a period of switching among multiple micro-states (or sub-states). Here, we will refer to a brain state as a sub-state during resting-state period. Of note, this multistablity perspective differs from studies on functional connectivity dynamics^15-23^, which have described the dynamic nature of the brain in terms of temporal changes in its interactions (connectivity parameters). In contrast, multistability in the complex system is an emergent property of nonlinear interactions among nodes in the system without any changes in their connectivity.

In a multistable system, stable states and the transition processes among them characterize the dynamics of the system. To explore the multistability and state transitions in the dynamic brain, energy landscape analysis has recently been applied to fMRI time series^24-30^. Prior to its introduction to the brain research field, energy landscape analysis had already shown its utility in understanding the dynamics of multi-dimensional complex systems, such as protein dynamics and the thermodynamics of liquids^31-36^. In the studies of brain dynamics using energy landscape analysis, distributed activity patterns across brain regions have often been used to define brain states, one of which the brain belongs to at each measurement time point^24-30^. In the energy landscape analysis, the energy of a state is the negative log probability of the occurrence of the state (thus, frequent states have low energy) according to the Boltzmann distribution of the state. The (inverse) frequency distribution of all possible brain states (patterns of brain activities across the brain regions) is called an energy landscape (see Figure 1).

**Figure 1.**
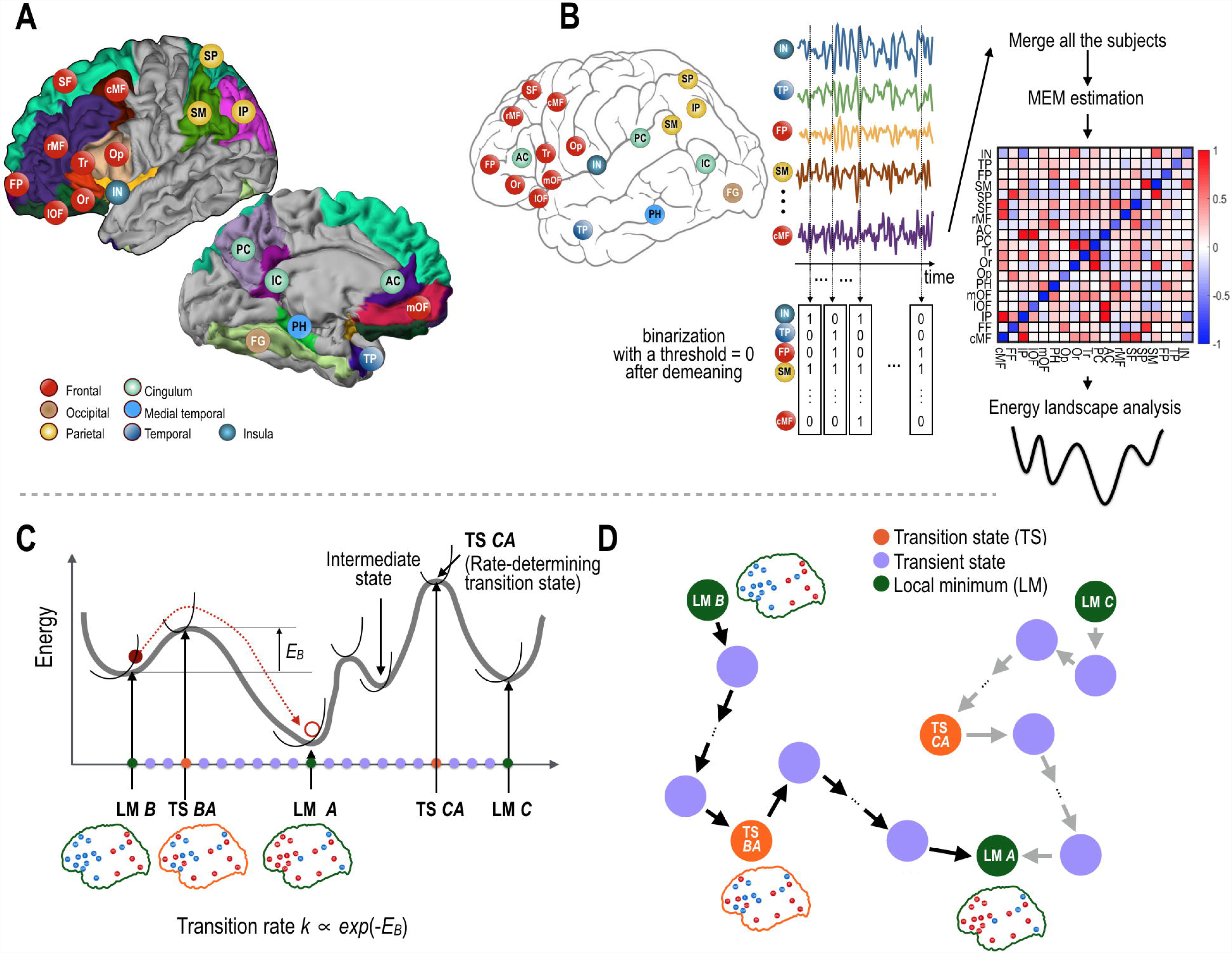
Procedures of the present study. (A) Regions of interest (ROIs) in the human cerebral cortex are shown. (B) Functional magnetic resonance (fMRI) data of the resting-state were binarized to represent brain states (active or inactive). These binarized states were used to construct the pairwise maximum entropy model, which was used to construct the energy landscape. (C, D) An illustration of the construction of the state transition network is presented. Local minima (LM) and transition states (saddle points, TS) on the transition pathway in the energy landscape (C), were used as nodes in the transition network as shown in (D). For each pathway, a transition rate was assigned as the weight on its edges (steps on the pathway) of the state transition network. Among transition states along the path, the transition state with the highest energy on the pathway determines the transition rate (called rate-determining transition state). Therefore, for simplicity, we refer to the rate-determining transition state as the TS for the transition between two states. LM A, LM B, LM C indicate local minima and TS BA indicates the transition from B to A, and TS CA, the transition from C to A. E_B_ indicates the energy barrier in the transition path. The red, green, and magenta color nodes represent local minima, transient, and transition state nodes, respectively.

The energy landscape of the system consists of several valleys with local minima (called “stable states” or “attractors”, abbreviated as LM) that have energies lower (more frequent) than their neighbors do in the valleys. Thus, the dynamics of the system can be divided into intravalley (within the basin of a local minimum) and intervalley (between-local minima) motions. In the former case, a state of the system wanders around a local minimum of the energy surface that the state belongs to, whereas in the latter case, a state transits from one local minimum to another, surpassing an energy barrier.

For a state transition between two local minima states, an optimal pathway refers to the path with the lowest maximal energy barrier among all possible paths. The optimal path may contain “intermediate states” (a type of local minima) and “transition states” (saddle point states) along the path. Among many transition states along the path, the transition state with the highest energy on the pathway determines the transition rate. Therefore, for brevity, we refer to this rate-determining transition state (having the highest energy on the pathway) as the transition state (TS) between two states (See Figure 1). Table 1 summarizes the terminology used in the current paper.

**Table 1.** Definitions of terminologies

| Terminology | Definition in this study |
| --- | --- |
| State, $V_i$ | A distributed brain activity pattern (a 19-dimensional binarized vector) |
| Stable state | A local minimum in the energy landscape, a brain activity pattern that sustains itself, and returns to its state following little perturbations. A state $V_i$ is a stable state, if it satisfies, $E(V_i) < E(V_{nei})$, where $V_{nei}$ are all neighbor states and $E(V_i)$ is the energy of the state $V_i$. |
| Neighbor state | A states $V_j$ is a neighbor state for a state $V_i$, if $d(V_i, V_j) = 1$, where $d(V_i, V_j)$ represents Hamming distance, i.e., the number of different elements (regional activity) in the vectors of two brain states $V_i$ and $V_j$. |
| Hub state | A highly connected (high node degree) state in the state transition network, indicating frequently occurring (or visiting) states while transitioning among different brain states. |
| Rate-determining transition state (TS) | When a transition path contains more than one local minima on the path (except for initial and final local minima), the transition path has multiple transition states (saddle points). The highest energy indicates the least occurrence. Among multiple transition states, the state with the highest energy determines the transition rate of transitioning between two states. The rate-determining transition state is referred to the transition state (TS) of the transition path between two states in the current study. |
| Intermediate state | A stable state on a transition path from a state to the other state, except for the initial and final states of the transition path. |
| Energy | Minus log (occurrence) probability of a state. Probability of appearance at a low energy state is higher than those of high energy states. In this study, the probability distribution of the states follows the Boltzmann distribution, which is given by Eq. 5. |
| Transition path | The path that has the minimal energy barrier among all possible paths from a local minimum to the other local minimum. |
| Transition rate | Transition speed from a local minimum to the other local minimum, defined as $\exp(-E_B)$ based on the state transition theory. $E_B$ is the energy barrier which is the energy difference between the highest energy on the transition path and the energy of the initial state. |
| STN | State transition network (STN) composed of brain states as nodes and transition rates as edges. |
| STN-FS | A full state transition network composed of all possible states along transition pathways between pairs of all stable states (local minima). |
| STN-GM | A state transition network composed of all states on the path from all local minima toward the global minimum. |
| STN-LM | A state transition network composed of rate-determining transition states and local minima. |

Despite a growing number of studies on the multistability of the resting brain systems^1,3-6,24,37-40^, the state transition processes between local minima in the brain systems have not yet been sufficiently investigated. In the present study, we explored the multistability and state transitional properties of the human cerebral cortex system. We estimated an energy landscape of brain states using a pairwise maximum entropy model (MEM) of the resting-state fMRI (rs-fMRI) data from the Human Connectome Project (HCP) database^41^. We extracted local minima and optimal pathways among them in the energy landscape, and then explored the characteristics of the brain state transition processes from the network-theoretical perspective using the state transition network, where states are represented as nodes, transitions between two states (nodes) as edges, and transition rates as edge weights.

In order to analyze the brain state transition, three (a full and two reduced) types of state transition networks were employed: a full state transition network composed of all possible states along transition pathways between pairs of all stable states (local minima) (STN-FS, Figure 2A), a state transition network composed of all states on the path from all local minima toward the global minimum (the most frequent brain state, STN-GM, Figure 2B), and a state transition network composed of all local minima and TSs (STN-LM, Figure 3A). In order to explore a general architecture of brain state transitions, we analyzed the STN-FS of brain dynamics with respect to the network theory using the degree of state nodes (occurrence frequency during state transition) and path lengths (how many transient states are needed to arrive at a final state). From this transition network analysis, we tested whether hub-like TSs exist, similar to spatial hubs found in the conventional network analysis of the brain^42-45^. We then narrowed down the STN-FS to the STN-GM to focus on state transition processes from local minima toward the global minimum. In the STN-GM analysis, we particularly examined whether state transitions toward the global minimum were processed in a single step or in multiple steps. If multi-step transitions existed, we then identified intermediate stable states (a type of local minima) that mediate those multi-step transitions. Finally, according to the transition state theory, a transition rate between two states is determined solely by the energy difference of the initial state and the rate-determining transition state (saddle point), i.e., the TS (Figure 1C). In order to explore the transition network in terms of the transition rate, we constructed the STN-LM using TSs and local minima. The hierarchy in the brain state transition was investigated by clustering the STN-LM with regard to the transition rate.

**Figure 2.**
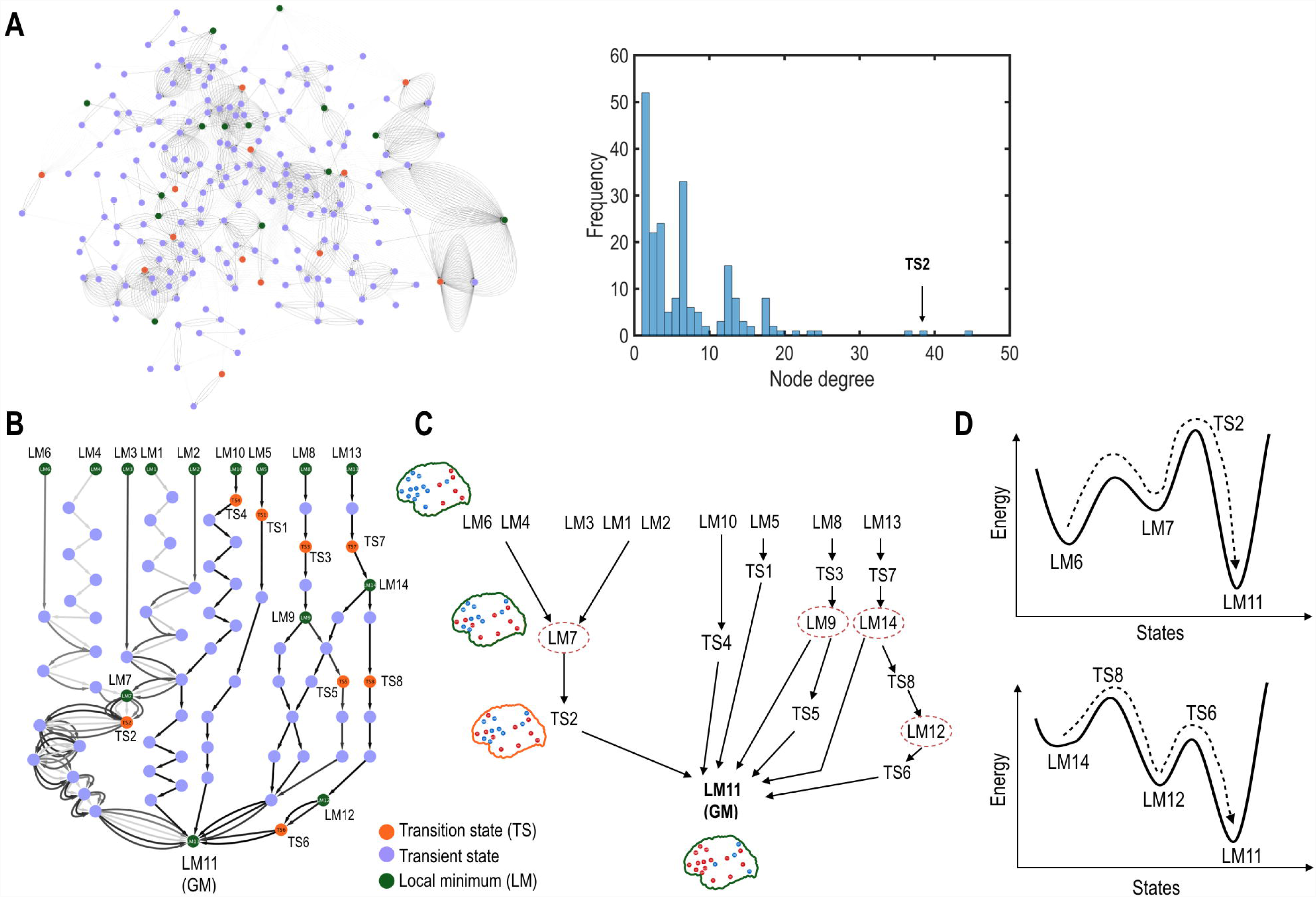
Analysis of the state transition networks. (A) The state transition network among full states (STN-FS) is shown in the left panel. We assigned all states in the state transition process to the nodes. The right panel shows a histogram of node degrees for all states (nodes) in the STN-FS. A transition state, TS2, has a significantly high node degree. (B - C) State transition processes (STN-GM) from local minima (LM) toward the global local minimum (LM11) is shown in (B). An illustration of the state transition processes in STN-GM is presented in (C). (D) Two representative examples, state transitions from LM6 to LM11 (upper panel) and from LM14 to LM11 (bottom panel), are shown. TS represents a transition state (a saddle point). The green, blue, and orange colors represent local minima, transient, and transition states, respectively.

**Figure 3.**
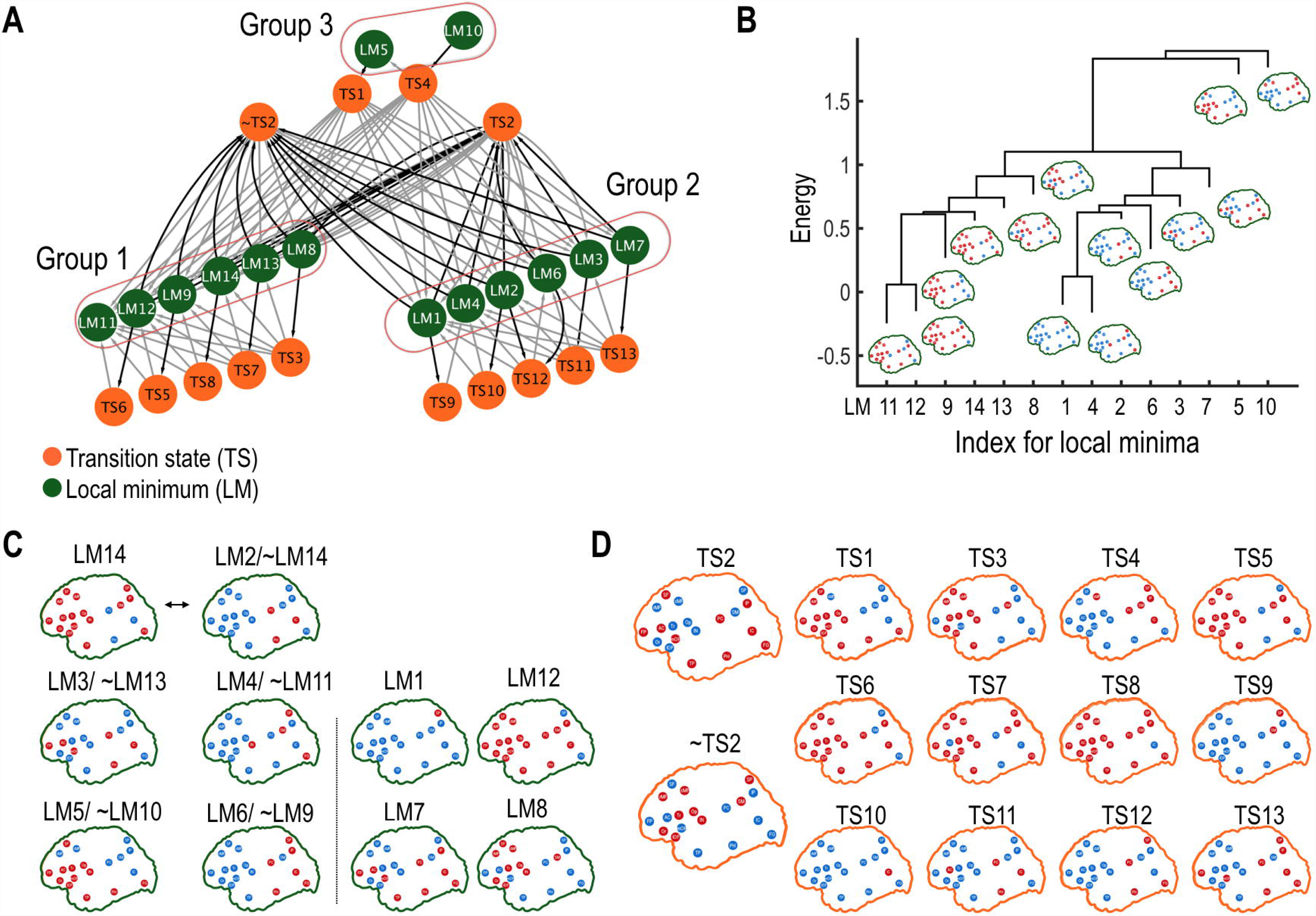
Analysis of the state transition network (STN-LM) composed of rate-determining transition states (TS) and local minima states (LM). (A) The STN-LM is shown. Black and gray colored lines represent an inward direction to and an outward direction from the TS. (B) Local minima (LM) were clustered according to energy barriers. The leaf ends of the dendrogram represent the energy values of the corresponding local minima. (C) Activation patterns of the local minima. The “∼” sign represents complementary states. For instance, LM2/∼LM14 indicates that LM2 and LM14 are each other’s complementary states. (D) Activity patterns of the transition states are shown with TS2 and ∼TS2 as major hub transition states. The red and blue dots represent the active and inactive states of the ROIs. The green and orange colors represent local minima and transition states.

We finally investigated the organizational properties of the resting-state brain by comparing the transitional properties of the baseline cortical system with those of a virtual system by altering the MEM parameters. The comparison between the resting-state brain and the altered (virtual) system was conducted over the STN-FS and STN-LM.

The results of our analysis suggest that the cerebral cortex system at rest contains multiple stable states that are clustered into two major state groups. The transition between brain states across the two state groups was mediated by a frequent TS, which operated as a hub of the transition network. When we removed this hub state, which bridges most transition processes across two groups, between-group transitions occurred via an alternative TS, indicating redundancy in state transition. State transition in the brain appears to involve multi-step state transitions, with some stable states serving as intermediate states for the complete transition. We also found that the baseline cerebral cortex at rest shows a more complex and organized state transition network than those of artificially altered systems. This network approach to the state transition in the brain may provide a new framework for the brain exploration and become an effective tool for understanding healthy and abnormal brain systems, concerning brain state dynamics.

## RESULTS

### Maximum entropy model for the cerebral cortex system at rest

In order to generate the energy landscape of the brain state, we estimated the first and second order interaction parameters (i.e., baseline sensitivity *H*_*i*_ and pairwise interaction *J*_*ij*_) of the MEM using binarized rs-fMRI activation patterns. The activation patterns of rs-fMRI data were reproduced with a high accuracy of fit (*r*_*D*_ = 86.3 %) and reliability (*ER* = 99.9 %) (Figure S1A). Baseline sensitivity parameters *H*_*i*_ and pairwise interaction, *J*_*ij*_, are displayed in Figure S1B and Figure 1B. Details of the obtained MEM parameters are described in supporting information (Figure S1).

### Multiple stable states in the resting-state of the cerebral cortex system

Analysis of the energy landscape identified 14 local minima of the cerebral cortex system at rest (Figures 2 and 3). Complementary states (active versus inactive for each brain region) of the five local minima were also found to be local minima (Figure 3). Two pairs, local minimum (LM) 1 and LM12, LM7, and LM8, were nearly complementary states of each other. In these pairs, all regions were complementary except for one brain region, in each: the inactive precuneus (PC) in the LM7 and LM8, and inactive fusiform gyrus (FG) in the LM1 and LM12.

The most stable local minimum (i.e., global minimum) was LM11, where most cortical regions were active except for the insula, supramarginal gyrus, superior parietal lobe, and fusiform gyrus (see LM11 map in Figure 2C).

### The state transition network among full states (STN-FS)

Utilizing disconnectivity graph analysis^57^, 91 transition pathways were extracted for all possible pairs of the 14 local minima. All states on the 91 transition pathways were regarded as nodes and transition rates between pairs of nodes as edges in the STN-FS (Figure 1C and 1D). As a result, a total of 219 nodes and 1201 edges composed an STN-FS (Figure 2A). When we evaluated node degrees for all nodes in the STN-FS, three (state) nodes showed a significantly higher node degree than the rest (Figure 2A). Most (86.8%) effective path lengths, i.e., the difference between the total path length and the Hamming distance of two initial and final local minima, were less than 8 (Figure S2). For half of the total number of pathways (49 pathways), effective path lengths had the shortest value of 1.

### Analysis of state transition network from local minima toward the global minimum (STN-GM)

For 14 local minima, 13 transition processes toward the global minimum were considered in the STN-GM with 82 nodes and 141 edges. A total of eight TSs, which determine transition rate, were found in the STN-GM. The STN-GM also showed that intermediate local minima (e.g., LM7, LM9, LM12, and LM14) were involved in the transition processes of other local minima transitioning toward the global minimum (Figures 2B and 2C). For instance, the transition pathways that started from the LM6, LM4, LM3, LM1, and LM12 passed through LM7 before reaching the global minimum (LM11). The rates for these transitions were determined by energy differences between TS2 and the initial local minima. The state transition from LM6 to the global minimum (LM11) contained an intermediate state (LM7) and the energy of the rate-determining transition state between LM6 and LM7 was smaller than that of LM7 and LM11 (i.e., energy of TS2), and, thus, the rate-determining transition state was TS2 (upper Figure 2D).

However, LM10, LM5, LM8, LM13, LM9, LM14, and LM12 had their own rate-determining transition states along transition paths toward the global minimum (Figures 2B and 2C). Indeed, in the state transition from LM14 to the global minimum (LM11), which contained an intermediate state (LM12), the energy of the rate-determining transition state between LM11 and LM12 (energy of TS8) was larger than that of LM12 and LM11 (energy of TS6), and, thus, the rate-determining transition state was TS8 (lower Figure 2D).

In this way, by analyzing this reduced state transition network (STN-GM), we could identify the characteristics of all transition processes on their way to the global minimum.

### Analysis of a state transition network among rate-determining transition states and local minima states (STN-LM)

The STN-LM was composed of 27 nodes (13 TSs plus 14 local minima) and 90 edges (Figure 3A). We found a clustered structure in the STN-LM: one cluster containing six local minima (LM11, LM12, LM9, LM14, LM13, and LM8), and the other containing their complementary local minima (LM1, LM4, LM2, LM6, LM3, and LM7). A similar clustering result was found by using energy barriers (Figure 3B). Interestingly, only one rate-determining transition state, TS2, bridged two clusters. TS2 is composed of active regions in the FP, SF, AC, mOF, PC, IP, IC, TP, PH, FG, which overlap mostly with coactivation of the default mode network^51^ and the anterior and medial temporal lobe (Figure 3D). Since TS2 has a high node degree (Figure 2A), we can refer to TS2 as a hub in the transition network.

To investigate the effects of TS2 on the transition process, we removed TS2 and explored the state transition pathways. After the removal of TS2, we found 36 alternative pathways. In these 36 pathways, the complementary state of TS2, namely ∼TS2, bridged the state transition processes between clusters, instead of TS2. The energy difference between TS2 and ∼TS2 was very small, 0.00737. Since the transition rate is proportional to *exp*(-*E*_barrier_) in state transition theory, the estimated ratio of the transition rate between the original and alternative pathway was 99.27%. Thus, we added the ∼TS2 node to the state transition network, STN-LM. Moreover, the property of clustered transitions was also observed for the transition processes in the ∼TS2 system (Figure S3A). Since TS2 and ∼TS2 have high node degrees (i.e., measure of frequency of appearance), both TS2 and ∼TS2 play as hub TSs.

We extracted the factors that determined transition rates (i.e., energy barriers) for both the TS2 system and TS2 removed (∼TS2) system (Figure 4); the Hamming distances between initial and final states were positively correlated with energy barriers (r = 0.608, p = 1.654 × 10^−10^ for the TS2 system, and r = 0.611 p = 1.276 ×10^−10^ for the ∼TS2 system, Figure 4B). However, there was no such relation between the energy barriers and effective path lengths (Figure S3C).

**Figure 4.**
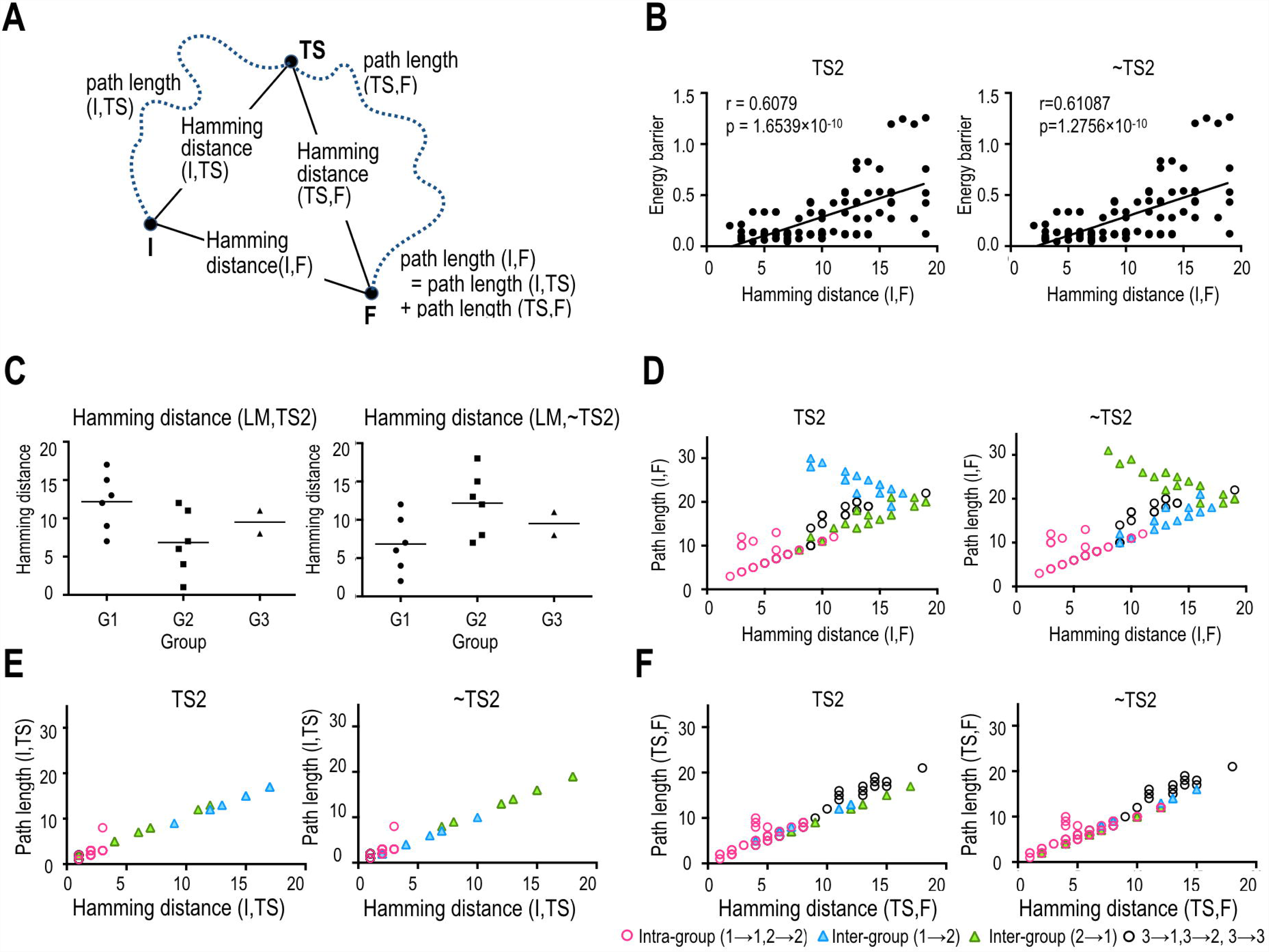
Path lengths and Hamming distances of the state transitions in the TS2 and ∼TS2 systems. The ∼TS2 system indicates transition pathways when the TS2 node was removed. (A) Definitions of the path lengths from the initial (I) to the final state (F) with transition state (TS) are presented. (B) Positive correlations between the energy barriers and Hamming distances of the initial and final states are presented. (C) Hamming distances between the local minima and TS2 (or ∼TS2) are presented for each group. (D) Full path lengths are plotted according to Hamming distances of the initial and final states. (E, F) Path lengths between initial and transition states (E), and path lengths between final and transition states (F) are plotted according to their Hamming distances. The definitions of the groups (G1, G2, and G3) are presented in Figure 3A. In (B) and (D) - (F), the left and right panels represent the results of the TS2 and ∼TS2 systems. The red points represent intra-group state transitions (i.e., transitions of group 1 → 1 and group 2 → 2). The blue and green triangle points represent the results of the state transitions between the inter-groups; group 1→2, (blue) and group 2→1 (green).

We further investigated the transition processes by separating the inter-group and intra-group processes (Figures 4D, 4E, and 4F). For the intra-group transitions, positive correlations between Hamming distances and path lengths were observed for both TS2 and ∼TS2 systems.

However, for the inter-group transitions, we could not find such associations. Thus, we further separated the inter-group transitions and found negative and positive correlations between Hamming distances and path lengths for the transitions from group 1 to 2 (r = −0.865, p = 3.123 × 10^−5^), and from group 2 to 1 (r = 0.921, p = 3.240 × 10^−9^) in the TS2 system (Figure 4D).

Interestingly, in the ∼TS2 system, the correlations were reversed; positive and negative correlations were found for the transitions from group 1 to 2 (r = 0.865, p = 3.123 × 10^−5^), and from group 2 to 1 (r = −0.921, p = 3.240 × 10^−9^) (Figure 4D). Since the distances between TS2 and local minima of group 1 were longer than those of group 2 in the TS2 system, but in the ∼TS2 system, an inverse association was found (i.e., the distances between ∼TS2 and local minima of group 1 were shorter than those of group 2) (Figure 4C), the cause of the inverse correlations could be the distance between the transition state and initial states. In the inter-group transitions, path lengths from an initial local minimum to the other group local minima depended on Hamming distances from the initial local minimum to the hub TS (Figure 4E and 4F).

### Effects on the state transition processes following the alteration of the system

The effects of the global strengths of pairwise interactions on the transition process were investigated by scaling all *J*_*ij*_ parameters (*αJ*_*ij*_, *α* = 0.0, 0.1, …, and 5.0). Both increases and decreases of the scale factor *α* tended to reduce the total number of local minima (Figure 5A). When a markedly small or large-scale factor, *α* < 0.7 or *α* > 1.7, was used, only one local minimum was found.

**Figure 5.**
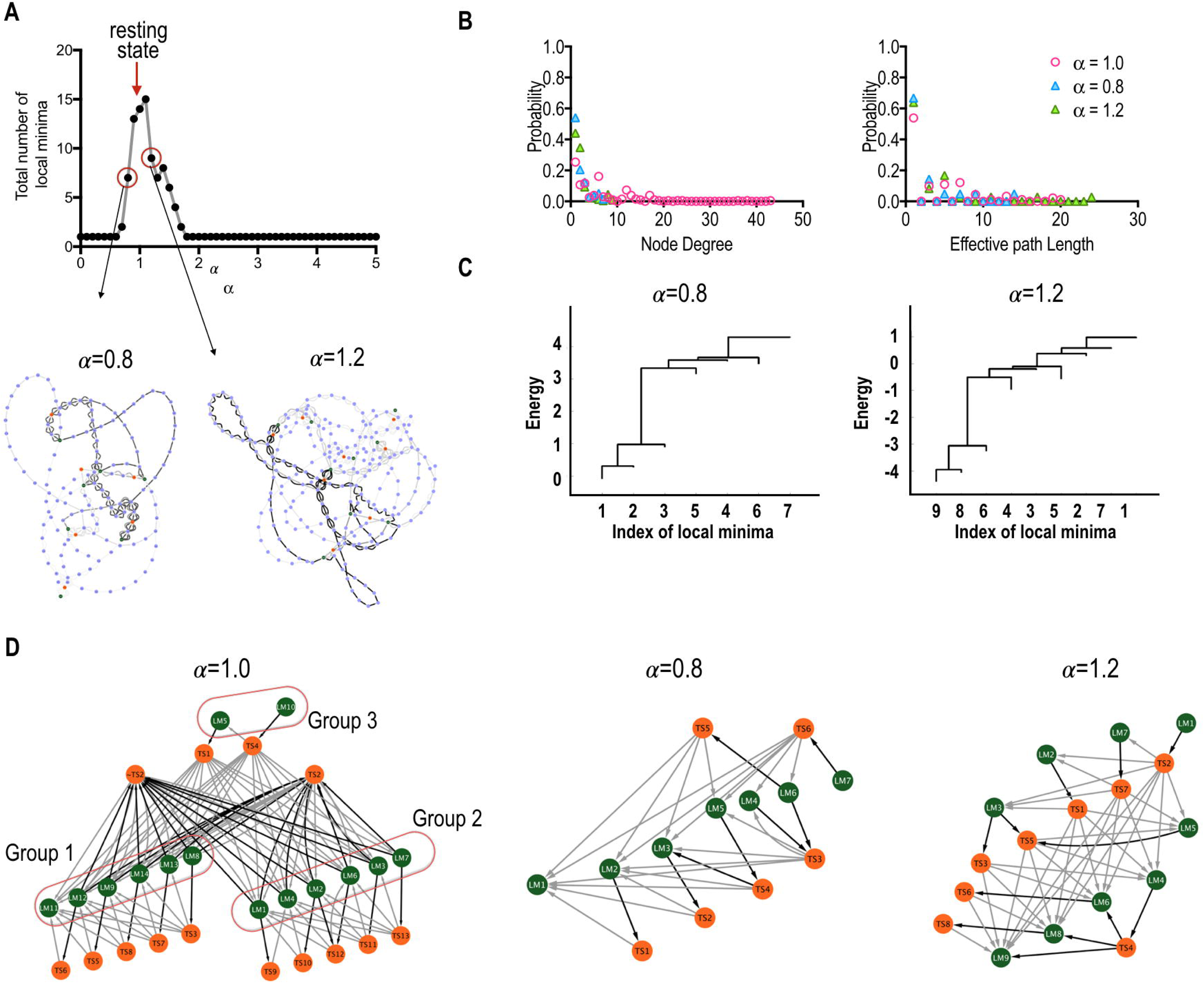
Comparisons between baseline and perturbed systems. (A) Total number of local minima in the perturbed systems, perturbed by changing global-scale pairwise interactions; αJ_ij_. The state transition networks among full states (STN-FS) of two representative cases (*α* = 0.8 and 1.2) are shown in the lower panel. (B) Probabilities of the node degree and effective path length in the state transition network are plotted. The red, blue, and green colors represent *α* = 1.0, *α* = 0.8 and 1.2. (C) Local minima were clustered according to energy barriers. The leaf ends of the dendrogram represent the energy values of the corresponding local minima. (D) The state transition networks (STNs-LM) composed of rate-determining transition states (TS) and local minima states (LM) of two perturbed systems are presented. The green and orange colored nodes in the reduced state transition networks represent local minima and transition states, respectively.

Here, we further analyzed two representative examples: *α* = 0.8 and *α* = 1.2. In both cases, the total number of local minima was reduced by perturbations, from 14 (baseline resting-state) to 7 (for *α* = 0.8) and 9 (for *α* = 1.2), and thus 21 and 36 transition processes were considered in their state transition networks, respectively.

In the state transition network of the weak coupling system (*α* = 0.8), 137 nodes and 258 edges, which were decreased compared to the baseline resting-state, were found. Under the strong pairwise interaction (*α* = 1.2), compared to the baseline resting-state, the total number of nodes was decreased (225 nodes) and that of edges was increased (456 edges). Since most pairwise parameters were positive, the energy of the states increased and decreased for weak and strong alterations in the scale parameter, respectively. In both cases, a positive correlation between node degree and energy was found for the nodes of the transient and transition states (Figure S4a).

In the baseline resting-state, nodes were densely connected to other nodes (Figure S4b), and, the maximum node degree was 44, which was larger than that of the altered systems (8 and 10, for *α* = 0.8, and 1.2, respectively). For all cases, more than half of the effective path lengths had a value of 1 (Figure 4). In the weak coupling system (*α* = 0.8), the longest effective path length was 15, which was smaller than that of the others (21 and 25 for *α* = 1.0, and 1.2, respectively).

In contrast to the baseline resting-state, in these altered systems, simple and deep energy valleys were found (Figure 5C). Indeed, the state transition processes were simpler than those of the baseline resting-state (Figure 5D). Except for LM6 in the weak coupling system, all local minima directly transitioned to their global minimum (Figure S4D).

## DISCUSSION

The brain at rest has been considered a highly dynamic complex system operating at a critical value of coupling that maximizes multistability^24,37-39,58^. Beyond the multistability of the resting-state cortical system, we systematically investigated the architecture of the state transition processes by applying a graph-theoretic analysis to state transition. State transition network analysis suggests a well-organized state transition process embedded in the resting-state human cerebral cortex system. The characteristics of the state transition in the resting state cortex system are discussed in the subsequent paragraphs.

The resting state brain has intermediate states in state dynamics. Some state transitions in the cerebral cortex system toward the global local minimum occurred in multi-steps via several intermediate stable states (or intermediate local minima) (Figure 2C and 6B). This phenomenon is similarly found in biochemical reactions by enzymes in biological systems^33,59-62^. For instance, during the rebinding of ligand (CO or O_2_ molecules) to myoglobin after photolysis, several intermediate states were observed in spectroscopic experiments, and these intermediates have often been explained in terms of the regulation of ligand binding mechanisms^60-62^. The state transitions during membrane fusion processes occur via intermediate states, which were explored in computational and experimental studies^63-65^. Similar to these phenomena in biochemical systems, the intermediate stable states during brain state transition may also play a role in lowering energy barriers. We speculate that this lowered energy barrier may regulate and expedite transitions along certain pathways of brain state transitions in the resting-state whole brain system. It should also be noted that transitions between some pairs of local minima are straightforward without any intermediate transition states.

Current network analysis of stable states of the cerebral cortex suggests several characteristics of brain state dynamics.

First, in the brain state transition network, local minima are highly clustered mainly into two groups, and the manner in which state transitions among distributed local minimum occurred was different between inter-group and intra-group transitions.

Second, most inter-group state transitions (from a local minimum at a cluster to a local minimum at the other) occurred via some hub transition states (saddle points) (e.g., TS2 and ∼TS2) in the transition pathway (Figure 6D). This phenomenon makes the inter-group transition different from the intra-group state transition, where the transition state along the path between two states differed according to the initial state. Those hub transition states are analogous to hubs found in the conventional network analysis of the brain connectome. Brain connectome studies have shown a hub-like structural architecture in the brain, which is considered to mediate efficient information exchange^42-45^. Similar to network analysis focusing on the spatial geometry of the connectome, the current result suggests that inter-group brain state transitions occur mostly via hub states (more frequently occurring states) in the temporal geometry (Figure 6C).

**Figure 6.**
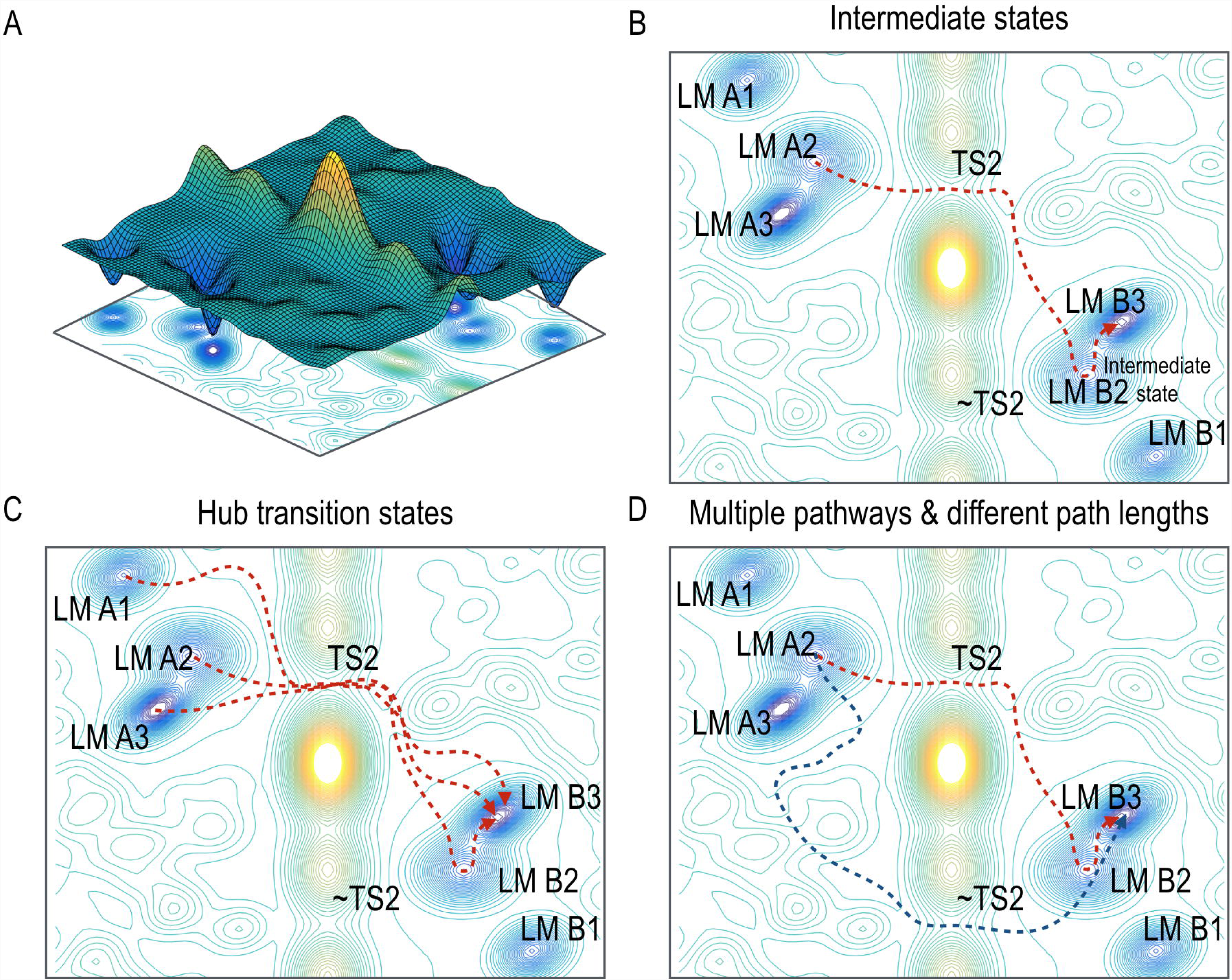
Illustrative analogy of the state transitions in the cerebral cortex system at rest. (A) A schematic energy landscape contains six stable states, which are classified into two major groups. (B) A representative pathway of the state transition is shown. Several stable states (local minima) operated as intermediate states of the state transition processes. (C) The hub transition state (saddle point) TS2 mediates the inter-group transition process between stable states in the two groups. (D) ∼TS2 also operated as an alternative hub transition state (saddle points) when TS2 was removed. This suggests multiple pathways and indicates redundant mechanisms of state transitions. Since the energy difference between the TS2 and ∼TS2 is small, transition rates of the two pathways, colored red and blue, are similar.

Third, path lengths (number of transitions to reach the target state) were positively correlated with the Hamming distances (i.e., number of different state bits (or regions)) between the initial and final states within the intra-group transitions. This result implies that the transition states of intra-group transitions take advantage of shorter transition paths, i.e., efficient transition from a brain state to the other state with minimal transition numbers. In the inter-group state transitions, path lengths were determined by the Hamming distance between the initial state and the hub transition state on the path, not the final state (Figures 4D, 4E, 4F, and 6D). We speculated that for significant state changes in the cortical system, the brain may minimize transitioning costs by traveling via hub transition states, not simply following short transition paths.

The current simulation study suggests that the cerebral cortex has redundant transition pathways. A transition state (e.g., TS2) mediates most of the inter-group state transition processes, serving as a transition hub in the resting-state transition network (STN-LM). When we excluded this hub transition state, its complementary state ∼TS2 appeared to serve as a detour for inter-group transitions with similar transition rates. This alternative hub ∼TS2 was the complementary state of the hub state (TS2) of the baseline resting-state. The energy level of ∼TS2 was similar to that of TS2 and the rates of the transition processes were similar to each other. We considered pathways via ∼TS2 as “redundant” pathway in inter-group transition processes, as those were chosen after removing TS2 as a transition state.

The existence of multiple pathways has been reported in some complex systems where a reaction occurs among multiple units cooperatively. For example, in some biomolecule systems, e.g. F1-ATPase and myosin V motor, two reaction pathways have been reported^66,67^. Multiple transition pathways may be associated with “degeneracy” or “redundancy” in the complex brain system^68,69^. The redundant pathways could be particularly advantageous in maintaining effective state transitions when a certain transient state cannot play its role in the state transition.

To understand the organization principle of the baseline configuration for brain state dynamics as done in Kang, et al. ^24^, we compared the baseline (observed) network configuration with the virtual (artificially altered) network configurations by scaling the pairwise interactions, and analyzed their state transition dynamics. Compared to the virtual networks, the baseline resting-state transition network (STN-FS) contained a bigger number of states with high node degrees and relatively longer path lengths. Furthermore, neither the clustered structure nor the intermediate states of the baseline system were observed in the altered virtual system. For example, in the STN-LM of the baseline system, which contains nodes on the pathway toward the global minimum (LM11) from other local minima (Figure 2), four types of transition processes were identified with four local minima (LM9, LM12, LM14, and LM7) as intermediate states. This property was not found in the virtual system, which showed a much simpler transition network (Figures 2 and S3).

Furthermore, hub intermediate states and hub transition states were found only in the baseline resting-state system, not in the virtual system. In the virtual systems, altered by scaling pairwise interactions from the baseline resting-state system, local minima having a high node degree disappeared; the hub-like local minima of the baseline system (having relatively high energy) were eliminated first by scaling pairwise interactions. This phenomenon, of which higher energy was eliminated first by network alteration, was consistent with that reported in our previous study on the subcortical system^24^. Meanwhile, low-energy local minima tended to persist even after network alteration. If we consider some task performances as states deviating from the baseline system, those sustaining local minima may act as common bases from which diverse functions arise or as fundamental elements of maintenance of dynamic brain systems. It may be that the baseline cerebral cortex system is configured to allow network systems to effectively transition among diverse brain states, which may be a necessary element in the workings of the complex network systems exhibiting multiple stable states.

All these transition characteristics may possibly be embedded in the nonlinear coupling over the structural network. We speculate that network topology may provide a biased playground of multistability, and endogenous fluctuation during resting-state may drive state transitions over the structural playground. This interpretation about the interplay between network topology and noise is in line with the dynamic nature of the brain^9,24^. In the current study, we showed that the multistable nature of brain states and the well-organized properties of the transition processes can emerge from nonlinear interactions over the cortical brain network.

Resting-state brain dynamics and non-stationary functional connectivity have recently been explored by evaluating brain-connectivity over the sliding windows from the viewpoint of a linear system^15-23^. For example, Park, et al. ^16^ assumed linear interactions (connectivity) between brain regions change by time, and these time-dependent interactions were estimated for consecutive windows. In contrast, multistability in the complex system is an emergent property of nonlinear interactions among nodes in the system, without any changes in the internal connectivity. From the perspective of the nonlinear system, Hansen, et al. ^40^ suggested that non-stationary functional connectivity (FC) (particularly, rapid transitions switching between a few discrete FC states) can be explained by the non-linearity of the nodal activity that derives the structural brain system. Spiegler, et al. ^70^ attributed the nonstationary FC to the criticality of the nonlinear brain system embedded in the structural network topology. Similarly, Cabral, et al. ^58^ also showed that dynamic functional connectivity can emerge from a static structural connectivity with various non-linear dynamic models of the brain. They showed that diverse FC states are emergent when the brain is operating at the edge of criticality. Pillai and Jirsa ^71^ also showed that multiple sub-states undergo structured flows on the manifold of the low-dimensional state spaces (functional subspaces) and this emergent behavior is attributable to the synaptic coupling level over the nonlinear interactions. Rabinovich, et al. ^72^ argues that both flexible and reproducible transitions among multiple meta-stable states can emerge in the nonlinear system, which may explain state transitions in the decision-making process. All those studies^40,58,70-72^ are based on a model with nonlinear temporal dynamics described using a differential equation. In terms of the nonlinear interaction and its consequent emergence of multiple states in the complex brain, our approach using pairwise MEM is in line with those studies^40,58,70-72^. However, the current evaluation differs from those studies in that the analysis of brain dynamics with a pairwise MEM is based on the statistical mechanics, which deals with the emergence of stable states and their transitions in terms of probability. Nevertheless, the two approaches (analysis with a differential equation and analysis in the statistical mechanics) are known to be equivalent since ensemble probability distributions of each state (in statistical mechanics) can be derived from a large number of trajectories as solutions of a differential equation (for a rigorous explanation, see ergodic theorem in the statistical physics).

A small number of major transition paths in the current study may serve as manifold-like transitions found in Pillai and Jirsa ^71^, where a lower dimensional manifold of state transitions was induced by an asymmetric interaction (due to task). The organized state transitions explored in this study may be correspondent to reproducible transitions among multiple meta-states in Rabinovich, et al. ^72^.

The current study has several limitations and challenges. Due to the high computational cost and the requirement of a large sample size, we evaluated the dynamics of a reduced brain system at each hemisphere (particularly focusing on the left hemisphere) but did not evaluate those of the whole brain system. In spite of strong symmetry e.g., Kang, et al. ^24^, the interaction between two hemispheres and its effect on the dynamic system in the whole brain system remain to be explored. We speculated that the dynamic properties of the whole-brain nonlinear system would be much more complex. In spite of technical challenges, exploration of the state transition properties of the whole brain system with more precisely parcellated brain regions would greatly expand our understanding of the brain system.

In the preprocessing step, we chose to conduct a global regression as we did for the subcortical brain system in our previous study^24^, which showed that the global regression emphasizes properties of the state dynamics. Since the current study focuses on the dynamics of brain sub-states in an equilibrium, i.e., a period without statistical changes in the global properties, and since the sub-states were defined in terms of spatially distributed activity patterns, we considered the relative distribution of brain activity as more adequate in representing dynamic brain states than factors due to global fluctuations.

Previous studies have revealed alterations in the dynamics of networks associated with brain disorders such as schizophrenia^73,74^ and autism^75^. A growing number of studies are showing altered dynamics in other brain disorders as well. However, the dynamic properties in brain diseases have not been thoroughly researched. The current frameworks for dynamic brain states can be used to identify altered dynamic architectures in neuropsychiatric disorders. Research using clinical data will yield results that can validate the usefulness of the proposed approach.

## MATERIALS AND METHODS

### Resting-state fMRI data set

The pairwise MEM (explained in the following section) was generated using rs-fMRI data of 470 participants (192 males, 278 females, ages: 29.19 ± 3.51 years) from the HCP database^41^, which was used in our previous study^24^. Briefly, all data were sampled at TR= 0.72 s, during 4 runs, with 1200 time points per run. A time series of the first principal component scores (i.e., eigenvalues) after applying a principal component analysis to fMRI time series at all voxels within a region was extracted as a signal for each region. The effects of rigid motion and their derivatives were regressed out, followed by linear detrending and despiking of the extracted signals^46-49^. Although there is an ongoing debate concerning filtering and global regression, we regressed out global signal changes in the whole-brain mask to emphasize short-term spatial patterns in representing specific brain states. Indeed, the current analysis is based on the assumption of the resting-state being in an equilibrium, i.e., without long-term statistical (temporal) changes in the dynamic properties.

Since computational cost dramatically increases by degree of the freedom of the system (2^N^, N: number of nodes), we extracted the rs-fMRI time series of only 19 regions of interest (ROIs) out of 33 cortical regions defined in the automated labeling map^50^. In choosing 19 ROIs to define a cortical state, we included brain regions associated with the default mode network, the salience network, and higher cognitive brain areas^51^ and excluded primary/secondary sensory and motor cortical regions in the evaluation. We also confined ROIs to a hemisphere (particularly to the left hemisphere) since previous studies showed strong symmetric activities (e.g., symmetric independent components found in many previous studies, including Smith, et al. ^52^) and strong interhemispheric connectivity e.g., Kang, et al. ^24^. Co-active regions across hemispheres behave similar time courses and, thus, are considered to be less informative in defining diverse brain states.

The ROIs used in this study are the precuneus (PC), parahippocampal gyrus (PH), caudal middle frontal gyrus (cMF), fusiform gyrus (FG), inferior parietal lobe (IP), isthmus cingulate gyrus (IC), lateral orbitofrontal gyrus (lOF), medial orbitofrontal (mOF), pars-opercularis (Op), pars-orbitalis (Or), pars-triangularis (Tr), rostral anterior cingulate gyrus (AC), rostral middle frontal gyrus (rMF), superior frontal gyrus (SF), superior parietal gyrus (SP), supramarginal gyrus (SM), frontal pole (FP), temporal pole (TP), and insula (IN) in the left hemisphere (Figure 1A). The ROIs in the left-hemisphere were mainly evaluated and presented in the current study. However, we confirmed that similar results were obtained from the ROIs in the right-hemisphere (see Supporting Information, Section S2).

For each ROI, signals were thresholded to represent inactive (0) and active (1) states. Since the number of local minima was maximized when the threshold was zero in the empirical evaluation (unpresented) and in our previous analysis^24^, we selected zero as the threshold to binarize regional states after global regression (Figure 1B). A brain state was defined by merging all (binarized) 19 regional states into a state vector (the number of elements of a state vector is 19). Due to a high sample size demand to estimate brain states (for all 2^19^ possible states), we concatenated all brain state samples from four sessions of 470 participants into a group-level sample data set (the total number of state samples, 1200 samples × 4 sessions × 470 participants) and estimated parameters of the group-level pairwise MEM using the method described in the following section.

### Construction of pairwise maximum entropy model (MEM)

To analyze resting-state activity in the cerebral cortex, we utilized the pairwise MEM estimation approach described in previous studies (Figure 1B)^24-26^.

The estimation process consists of a step for defining brain states and a pairwise MEM model for state dynamics, and an optimization step for MEM model parameters to fit probability distributions of empirical brain states and states generated by the model. Details of the model construction are provided in our previous study^24^ (for the mathematical details, see review ref.^53^).

Briefly, the brain state at time *t* is represented as a state vector:

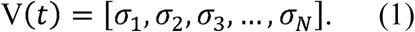

The value of *σ* _*i*_ is either 0 (inactive) or 1 (active), and *N* represents the total number of nodes (or ROIs).

In the pairwise MEM, the average of each node activity,

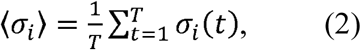

and the averages of all pairwise products,

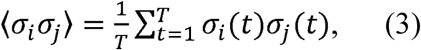

are fixed to characterize the system. With these constraints, maximizing the entropy *S*

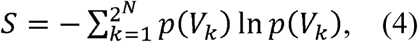

derives the following probability of the state *V*_*k*_, *p*(*V*_*k*_), as a Boltzmann distribution

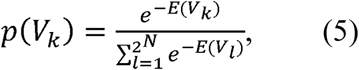

where *E*(*V*_*k*_) represents the energy of the state *V*_*k*_,

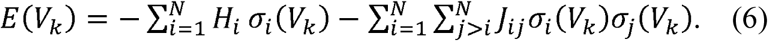

The parameters *H*_*i*_ and *J*_*ij*_ represent the activation tendency (baseline sensitivity) of node *i* and the pairwise interaction between nodes *i* and *j*, respectively. A gradient ascent algorithm was employed to estimate MEM parameters, *H*_*i*_ and *J*_*ij*_. These parameters were iteratively updated using the following equations,

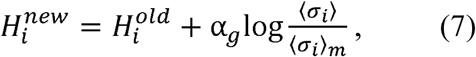

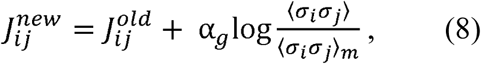

Here, ⟨*σ*_*i*_⟩_*m*_ and ⟨*σ*_*i*_ *σ*_*j*_⟩_*m*_ were calculated as follows:

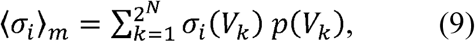

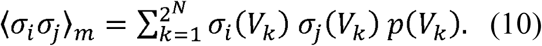

Scale parameter *a*_*g*_ was initially set to 0.1. The parameters were optimized until the gradients reached a value lower than 10^−5^. In the calculation for the experimental probability distribution of brain states, we calculated the frequency of each state in the group-level sample data set described above.

To evaluate the effectiveness of the pairwise MEM, we calculated the accuracy value, *r*_*D*_,

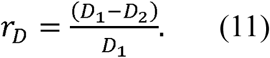

Here, *D*_*k*_ is the Kullback-Leibler divergence between the probability distributions of the *k*-th order model network and the empirical network,

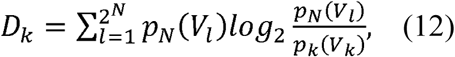

where *p*_*N*_ represents the empirical distribution of the network state. We also evaluated the reliability parameter *ER*,

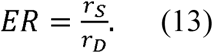

Here, *r*_*s*_ and *S*_*k*_ are given by

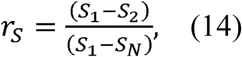

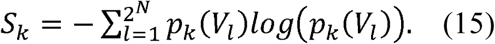

The measures *r*_*D*_ and *r*_*S*_ evaluate the adequacy of the pairwise MEM over the independent MEM in explaining time series, in two different aspects; *r*_*D*_ and *r*_*S*_ use Kullback-Leibler divergences and difference of the entropy between independent (1st order) and pairwise (2nd order) MEMs. The reliability *ER* was defined to compare those two different measures. If *H*_*i*_ and *J*_*ij*_ are estimated without error, *ER* is equal to 1.

### Energy landscape analysis

To describe the dynamics of the cerebral cortex system at rest, we performed the energy landscape analysis. More specifically, first, we elucidated the local minima (attractors), and then evaluated energy barriers between pairs of attractors, following the procedure described in the previous paper^24,26^.

To construct an energy landscape, the distance between two states should be first defined. Based on this distance, neighbor states can be defined to extract local minima. Following previous energy landscape studies, we defined the distance between two states as the number of elements (bits) that differ between two state vectors. We also assumed a gradual state transition, and the energy landscape was examined by changing one element of the state vector for each step.

The local minima (also called, stable states) were defined in states that have lower energy (more frequent) relative to their neighbors. To evaluate the energy barrier for each local minima pair, the lowest energy pathways were extracted by using the disconnectivity graph analysis^54^. Specifically, for each possible pair of local minima, we recorded the shortest path connecting the two local minima. The highest energy on this path was selected as a threshold to remove states that exhibited higher energy than the threshold. We repeated this step until the two local minima had been disconnected. The highest energy value of the last connected path was assigned to the threshold of the local minimum pair. The energy barrier, *E*_*B*_, between two local minima *i* and *j*, was defined as the lower value between *E*_*th*_(*V*_*i*_,*V*_*j*_) – *V*_*i*_ and *E*_*th*_ (*V*_*i*_,*V*_*j*_) – *V*_j_, where *E*_*th*_ (*V*_*i*_,*V*_*j*_) represents the threshold as defined above. These disconnectivity graph calculations were performed using the i-graph library^55^.

### Construction of the state transition networks

In the present study, we constructed three types of state transition networks; a state transition network composed of all possible states along transition pathways among local minima (STN-FS), a state transition network of states from local minima toward the global minimum (STN-GM), and a state transition network among TSs and local minima states (STN-LM).

We first constructed a STN-FS as a directional weighted network (Figure 2A). For all possible pairs of local minima, state transition pathways were identified as described in the above section. All states on the state transition pathways among local minima were considered nodes of the STN-FS. Since the forward and backward state transition pathways were identical for a pair of local minima, we only considered the state transitions from the higher to lower local minima. For all edges (transitions between pairs of states), we assigned weights using a transition rate, which is given by

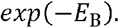

Here, *E*_*B*_ represents the energy barrier (or activation energy) which is the energy difference between the highest energy on the transition path and the energy of the initial state (see Figure 1C).

We then constructed the STN-GM to focus on the details of the state transition processes toward the global minimum (Figure 2B). To construct this network, all nodes and edges that were not connected to the global minimum in the STN-FS were removed. In the STN-GM analysis, we particularly identified intermediate states that mediate multi-step transitions toward the global minimum.

In order to focus on the transition rate among local minima, we constructed a STN-LM (Figure 3A), considering local minima and TSs as nodes and transition rates as edges. Note that the transition rate between two states only depended on its energy barrier (maximal energy difference) along the pathway. Based on the transition rates in STN-LM, we evaluated the architecture of the state transitions of the system by applying a cluster analysis of transition rates to differentiate intra- and inter-group transitions. The clustering of transitions in the STN-LM was conducted using the UPGMA (Unweighted Pair Group Method with Arithmetic Mean) algorithm^56^ with Euclidian metric; distances between nodes were defined by transition rates.

The constructed network was analyzed in terms of the network theory using node degree (frequency of occurrence during state transition) and path length (number of transitions to reach the target state). We also analyzed the effective path length between local minima, which is defined as the difference between the total path length (number of transitions) and the Hamming distance (number of different elements between two state vectors, i.e., number of regions that showed active/inactive differences) of the initial and final states.

## Acknowledgments

This research was supported by Brain Research Program and the Korea Research Fellowship Program through the National Research Foundation of Korea (NRF) funded by the Ministry of Science and ICT (NRF-2017M3C7A1049051 and NRF-2017H1D3A1A01053094). Data were provided in part by the Human Connectome Project, WU-Minn Consortium (Principal Investigators: David Van Essen and Kamil Ugurbil; 1U54MH091657) funded by the 16 NIH Institutes and Centers that support the NIH Blueprint for Neuroscience Research; and by the McDonnell Center for Systems Neuroscience at Washington University. The authors thank Ms. Hanseul Choi for her support in the writing.

## Author Contributions

J.K. and H.J.P. developed the method and wrote the manuscript. J.K. performed calculations and analyzed the results. C.P. participated in pre-processing of the fMRI data.

## Data availability

All fMRI data analyzed in the present study were obtained from the Human Connectome Project database, which is publicly accessible at: http://www.humanconnectome.org.

## References

1 Deco, G., Tononi, G., Boly, M. & Kringelbach, M. L. Rethinking segregation and integration: contributions of whole-brain modelling. Nature reviews. Neuroscience 16, 430–439, doi:10.1038/nrn3963 (2015).

2 Deco, G. & Jirsa, V. K. Ongoing cortical activity at rest: criticality, multistability, and ghost attractors. The Journal of neuroscience : the official journal of the Society for Neuroscience 32, 3366–3375, doi:10.1523/JNEUROSCI.2523-11.2012 (2012).

3 Cabral, J., Kringelbach, M. L. & Deco, G. Exploring the network dynamics underlying brain activity during rest. Progress in neurobiology 114, 102–131, doi:10.1016/j.pneurobio.2013.12.005 (2014).

4 Tognoli, E. & Kelso, J. A. The metastable brain. Neuron 81, 35–48, doi:10.1016/j.neuron.2013.12.022 (2014).

5 Freyer, F., Roberts, J. A., Ritter, P. & Breakspear, M. A canonical model of multistability and scale-invariance in biological systems. PLoS Comput Biol 8, e1002634, doi:10.1371/journal.pcbi.1002634 (2012).

6 Freyer, F. et al. Biophysical mechanisms of multistability in resting-state cortical rhythms. The Journal of neuroscience : the official journal of the Society for Neuroscience 31, 6353–6361, doi:10.1523/JNEUROSCI.6693-10.2011 (2011).

7 Kelso, J. A. Multistability and metastability: understanding dynamic coordination in the brain. Philos Trans R Soc Lond B Biol Sci 367, 906–918, doi:10.1098/rstb.2011.0351 (2012).

8 Schwartz, J. L., Grimault, N., Hupe, J. M., Moore, B. C. & Pressnitzer, D. Multistability in perception: binding sensory modalities, an overview. Philos Trans R Soc Lond B Biol Sci 367, 896–905, doi:10.1098/rstb.2011.0254 (2012).

9 Breakspear, M. Dynamic models of large-scale brain activity. Nature neuroscience 20, 340–352, doi:10.1038/nn.4497 (2017).

10 Rabinovich, M. I. & Varona, P. Robust transient dynamics and brain functions. Front Comput Neurosci 5, 24, doi:10.3389/fncom.2011.00024 (2011).

11 Laird, A. R. et al. Behavioral interpretations of intrinsic connectivity networks. Journal of cognitive neuroscience 23, 4022–4037, doi:10.1162/jocn_a_00077 (2011).

12 Smith, S. M. et al. Correspondence of the brain’s functional architecture during activation and rest. Proceedings of the National Academy of Sciences of the United States of America 106, 13040–13045, doi:10.1073/pnas.0905267106 (2009).

13 Damoiseaux, J. S. et al. Consistent resting-state networks across healthy subjects. Proceedings of the National Academy of Sciences of the United States of America 103, 13848–13853, doi:10.1073/pnas.0601417103 (2006).

14 Park, B., Kim, D. S. & Park, H. J. Graph independent component analysis reveals repertoires of intrinsic network components in the human brain. PLoS One 9, e82873, doi:10.1371/journal.pone.0082873 (2014).

15 Monti, R. P. et al. Estimating time-varying brain connectivity networks from functional MRI time series. NeuroImage 103, 427–443, doi:10.1016/j.neuroimage.2014.07.033 (2014).

16 Park, H.-J., Friston, K. J., Pae, C., Park, B. & Razi, A. Dynamic effective connectivity in resting state fMRI. NeuroImage, doi:https://doi.org/10.1016/j.neuroimage.2017.11.033 (2017).

17 Jeong, S. O., Pae, C. & Park, H. J. Connectivity-based change point detection for large-size functional networks. NeuroImage 143, 353–363, doi:10.1016/j.neuroimage.2016.09.019 (2016).

18 Hutchison, R. M., Womelsdorf, T., Gati, J. S., Everling, S. & Menon, R. S. Resting-state networks show dynamic functional connectivity in awake humans and anesthetized macaques. Hum Brain Mapp 34, 2154–2177, doi:10.1002/hbm.22058 (2013).

19 Handwerker, D. A., Roopchansingh, V., Gonzalez-Castillo, J. & Bandettini, P. A. Periodic changes in fMRI connectivity. NeuroImage 63, 1712–1719, doi:10.1016/j.neuroimage.2012.06.078 (2012).

20 Chang, C. & Glover, G. H. Time-frequency dynamics of resting-state brain connectivity measured with fMRI. NeuroImage 50, 81–98, doi:10.1016/j.neuroimage.2009.12.011 (2010).

21 Allen, E. A. et al. Tracking whole-brain connectivity dynamics in the resting state. Cereb Cortex 24, 663–676, doi:10.1093/cercor/bhs352 (2014).

22 Cribben, I., Haraldsdottir, R., Atlas, L. Y., Wager, T. D. & Lindquist, M. A. Dynamic connectivity regression: determining state-related changes in brain connectivity. NeuroImage 61, 907–920, doi:10.1016/j.neuroimage.2012.03.070 (2012).

23 Calhoun, V. D., Miller, R., Pearlson, G. & Adali, T. The chronnectome: time-varying connectivity networks as the next frontier in fMRI data discovery. Neuron 84, 262–274, doi:10.1016/j.neuron.2014.10.015 (2014).

24 Kang, J., Pae, C. & Park, H. J. Energy landscape analysis of the subcortical brain network unravels system properties beneath resting state dynamics. NeuroImage 149, 153–164, doi:10.1016/j.neuroimage.2017.01.075 (2017).

25 Watanabe, T. et al. A pairwise maximum entropy model accurately describes resting-state human brain networks. Nat Commun 4, 1370, doi:10.1038/ncomms2388 (2013).

26 Watanabe, T. et al. Energy landscapes of resting-state brain networks. Frontiers in neuroinformatics 8, 12, doi:10.3389/fninf.2014.00012 (2014).

27 Watanabe, T. et al. Network-dependent modulation of brain activity during sleep. NeuroImage 98, 1–10, doi:10.1016/j.neuroimage.2014.04.079 (2014).

28 Watanabe, T., Masuda, N., Megumi, F., Kanai, R. & Rees, G. Energy landscape and dynamics of brain activity during human bistable perception. Nat Commun 5, 4765, doi:10.1038/ncomms5765 (2014).

29 Ezaki, T., Sakaki, M., Watanabe, T. & Masuda, N. Age-related changes in the ease of dynamical transitions in human brain activity. Hum Brain Mapp 39, 2673–2688, doi:10.1002/hbm.24033 (2018).

30 Gu, S. et al. The Energy Landscape of Neurophysiological Activity Implicit in Brain Network Structure. Sci Rep 8, 2507, doi:10.1038/s41598-018-20123-8 (2018).

31 Rao, F. & Karplus, M. Protein dynamics investigated by inherent structure analysis. Proceedings of the National Academy of Sciences of the United States of America 107, 9152–9157, doi:10.1073/pnas.0915087107 (2010).

32 Li, C. B., Yang, H. & Komatsuzaki, T. Multiscale complex network of protein conformational fluctuations in single-molecule time series. Proceedings of the National Academy of Sciences of the United States of America 105, 536–541, doi:10.1073/pnas.0707378105 (2008).

33 Frauenfelder, H., Sligar, S. G. & Wolynes, P. G. The energy landscapes and motions of proteins. Science 254, 1598–1603 (1991).

34 Gfeller, D., De Los Rios, P., Caflisch, A. & Rao, F. Complex network analysis of free-energy landscapes. Proceedings of the National Academy of Sciences of the United States of America 104, 1817–1822, doi:10.1073/pnas.0608099104 (2007).

35 Delvenne, J. C., Yaliraki, S. N. & Barahona, M. Stability of graph communities across time scales. Proceedings of the National Academy of Sciences of the United States of America 107, 12755–12760, doi:10.1073/pnas.0903215107 (2010).

36 Goldstein, M. Viscous Liquids and the Glass Transition: A Potential Energy Barrier Picture. The Journal of Chemical Physics 51, 3728–3739, doi:10.1063/1.1672587 (1969).

37 Golos, M., Jirsa, V. & Dauce, E. Multistability in Large Scale Models of Brain Activity. PLoS Comput Biol 11, e1004644, doi:10.1371/journal.pcbi.1004644 (2015).

38 Deco, G., Senden, M. & Jirsa, V. How anatomy shapes dynamics: a semi-analytical study of the brain at rest by a simple spin model. Front Comput Neurosci 6, 68, doi:10.3389/fncom.2012.00068 (2012).

39 Deco, G., Jirsa, V. K. & McIntosh, A. R. Resting brains never rest: computational insights into potential cognitive architectures. Trends in neurosciences 36, 268–274, doi:10.1016/j.tins.2013.03.001 (2013).

40 Hansen, E. C., Battaglia, D., Spiegler, A., Deco, G. & Jirsa, V. K. Functional connectivity dynamics: modeling the switching behavior of the resting state. NeuroImage 105, 525–535, doi:10.1016/j.neuroimage.2014.11.001 (2015).

41 Van Essen, D. C. et al. The Human Connectome Project: a data acquisition perspective. NeuroImage 62, 2222–2231, doi:10.1016/j.neuroimage.2012.02.018 (2012).

42 Bassett, D. S. & Bullmore, E. T. Small-World Brain Networks Revisited. The Neuroscientist 26, 107385841666772–107385841666718 (2017).

43 Senden, M., Reuter, N., van den Heuvel, M. P., Goebel, R. & Deco, G. Cortical rich club regions can organize state-dependent functional network formation by engaging in oscillatory behavior. NeuroImage (2016).

44 van den Heuvel, M. P. & Sporns, O. Rich-club organization of the human connectome. The Journal of neuroscience : the official journal of the Society for Neuroscience 31, 15775–15786 (2011).

45 Honey, C. J. & Sporns, O. Dynamical consequences of lesions in cortical networks. Hum Brain Mapp 29, 802–809 (2008).

46 Weissenbacher, A. et al. Correlations and anticorrelations in resting-state functional connectivity MRI: a quantitative comparison of preprocessing strategies. NeuroImage 47, 1408–1416, doi:10.1016/j.neuroimage.2009.05.005 (2009).

47 Power, J. D., Barnes, K. A., Snyder, A. Z., Schlaggar, B. L. & Petersen, S. E. Spurious but systematic correlations in functional connectivity MRI networks arise from subject motion. NeuroImage 59, 2142–2154, doi:10.1016/j.neuroimage.2011.10.018 (2012).

48 Thomas, J. B. et al. Functional connectivity in autosomal dominant and late-onset Alzheimer disease. JAMA neurology 71, 1111–1122, doi:10.1001/jamaneurol.2014.1654 (2014).

49 Taylor, J. S., Rastle, K. & Davis, M. H. Interpreting response time effects in functional imaging studies. NeuroImage 99, 419–433, doi:10.1016/j.neuroimage.2014.05.073 (2014).

50 Desikan, R. S. et al. An automated labeling system for subdividing the human cerebral cortex on MRI scans into gyral based regions of interest. NeuroImage 31, 968–980, doi:10.1016/j.neuroimage.2006.01.021 (2006).

51 Power, J. D. et al. Functional network organization of the human brain. Neuron 72, 665–678, doi:10.1016/j.neuron.2011.09.006 (2011).

52 Smith, S. M. et al. Resting-state fMRI in the Human Connectome Project. NeuroImage 80, 144–168, doi:10.1016/j.neuroimage.2013.05.039 (2013).

53 Yeh, F. C. et al. Maximum Entropy Approaches to Living Neural Networks. Entropy 12, 89–106, doi:10.3390/e12010089 (2010).

54 Becker, O. M. & Karplus, M. The topology of multidimensional potential energy surfaces: Theory and application to peptide structure and kinetics. The Journal of Chemical Physics 106, 1495–1517, doi:10.1063/1.473299 (1997).

55 Csárdi, G. & Nepusz, T. The igraph software package for complex network research. Inter Journal Complex Systems, 1695 (2006).

56 Sokal, R. & Michener, C. A statistical method for evaluating systematic relationships. University of Kansas Science Bulletin 38, 1409–1438 (1958).

57 Becker, O. M. & Karplus, M. The topology of multidimensional potential energy surfaces: Theory and application to peptide structure and kinetics. J. Chem. Phys. 106, 1495–1517, doi:Doi 10.1063/1.473299 (1997).

58 Cabral, J., Kringelbach, M. L. & Deco, G. Functional connectivity dynamically evolves on multiple time-scales over a static structural connectome: Models and mechanisms. NeuroImage 160, 84–96, doi:10.1016/j.neuroimage.2017.03.045 (2017).

59 Stagno, J. R. et al. Structures of riboswitch RNA reaction states by mix-and-inject XFEL serial crystallography. Nature, doi:10.1038/nature20599 (2016).

60 Yuan, Y., Tam, M. F., Simplaceanu, V. & Ho, C. New look at hemoglobin allostery. Chem Rev 115, 1702–1724, doi:10.1021/cr500495x (2015).

61 Vesper, M. D. & de Groot, B. L. Collective dynamics underlying allosteric transitions in hemoglobin. PLoS Comput Biol 9, e1003232, doi:10.1371/journal.pcbi.1003232 (2013).

62 Mihailescu, M. R. & Russu, I. M. A signature of the T ---> R transition in human hemoglobin. Proceedings of the National Academy of Sciences of the United States of America 98, 3773–3777, doi:10.1073/pnas.071493598 (2001).

63 François-Martin, C., Rothman, J. E. & Pincet, F. Low energy cost for optimal speed and control of membrane fusion. Proceedings of the National Academy of Sciences of the United States of America 114, 1238–1241 (2017).

64 Ryham, R. J., Klotz, T. S., Yao, L. & Cohen, F. S. Calculating Transition Energy Barriers and Characterizing Activation States for Steps of Fusion. Biophysical journal 110, 1110– 1124 (2016).

65 Smirnova, Y. G., Marrink, S.-J., Lipowsky, R. & Knecht, V. Solvent-Exposed Tails as Prestalk Transition States for Membrane Fusion at Low Hydration. J. Am. Chem. Soc. 132, 6710–6718 (2010).

66 Shimabukuro, K., Muneyuki, E. & Yoshida, M. An alternative reaction pathway of F1- ATPase suggested by rotation without 80 degrees/40 degrees substeps of a sluggish mutant at low ATP. Biophys J 90, 1028–1032, doi:10.1529/biophysj.105.067298 (2006).

67 Uemura, S., Higuchi, H., Olivares, A. O., De La Cruz, E. M. & Ishiwata, S. Mechanochemical coupling of two substeps in a single myosin V motor. Nat Struct Mol Biol 11, 877–883, doi:10.1038/nsmb806 (2004).

68 Price, C. J. & Friston, K. J. Degeneracy and cognitive anatomy. Trends Cogn Sci 6, 416– 421 (2002).

69 Edelman, G. M. & Gally, J. A. Degeneracy and complexity in biological systems. Proceedings of the National Academy of Sciences of the United States of America 98, 13763–13768, doi:10.1073/pnas.231499798 (2001).

70 Spiegler, A., Hansen, E. C., Bernard, C., McIntosh, A. R. & Jirsa, V. K. Selective Activation of Resting-State Networks following Focal Stimulation in a Connectome-Based Network Model of the Human Brain. eNeuro 3, doi:10.1523/ENEURO.0068-16.2016 (2016).

71 Pillai, A. S. & Jirsa, V. K. Symmetry Breaking in Space-Time Hierarchies Shapes Brain Dynamics and Behavior. Neuron 94, 1010–1026, doi:10.1016/j.neuron.2017.05.013 (2017).

72 Rabinovich, M. I., Huerta, R., Varona, P. & Afraimovich, V. S. Transient cognitive dynamics, metastability, and decision making. PLoS Comput Biol 4, e1000072, doi:10.1371/journal.pcbi.1000072 (2008).

73 Loh, M., Rolls, E. T. & Deco, G. A dynamical systems hypothesis of schizophrenia. PLoS Comput Biol 3, e228 (2007).

74 Cabral, J. et al. Structural connectivity in schizophrenia and its impact on the dynamics of spontaneous functional networks. Chaos: An Interdisciplinary Journal of Nonlinear Science 23, 046111, doi:10.1063/1.4851117 (2013).

75 Watanabe, T. & Rees, G. Brain network dynamics in high-functioning individuals with autism. Nature Communications 8, 16048, doi:10.1038/ncomms16048 https://www.nature.com/articles/ncomms16048-supplementary-information (2017).

